# Structural heterogeneity of uncrosslinked FtsH-HflKC assemblies includes variable FtsH occupancy and a distinct wild-type FtsH architecture

**DOI:** 10.64898/2026.10.12.758600

**Authors:** Günce Göc, Deniz Ugurlar, Imre Berger, Christiane Schaffitzel, Burak V. Kabasakal

## Abstract

The AAA+ metalloprotease FtsH associates with HflK and HflC to form the megadalton FtsH-HflKC complex (FHC), which contributes to membrane-protein quality control in *Escherichia coli*. Recent structural studies have revealed asymmetric HflKC architectures and variable FtsH occupancy, contrasting with earlier crosslinked structures of a symmetric four-hexamer cage. Here, we analyzed wild-type, uncrosslinked FHC co-expressed from a single multicistronic plasmid and solubilized in lauryl maltose neopentyl glycol (LMNG) using single-particle cryo-EM, SEC-SAXS and biochemical activity assays. 2D classification identified HflKC-containing particles associated with one or two FtsH hexamers, together with a distinct population of isolated FtsH particles. Focused reconstruction of the FtsH cytosolic domain reached 5.32 Å resolution and provided a structural view of wild-type, uncrosslinked FtsH without stabilizing substitutions or an added nucleotide analogue. The reconstruction retained the characteristic stacked arrangement of the AAA+ and protease rings but showed a more laterally splayed AAA+ ring compared with the stabilized reference, while the protease ring remained broadly similar in overall envelope. A 9.63 Å HflKC consensus reconstruction and 3D variability analysis (3DVA) in C1, further revealed substantial conformational variability spanning open-like and more constricted scaffold geometries. SEC-SAXS of a selected later-eluting LMNG-solubilized fraction yielded a compact anisotropic envelope with an estimated *D*_max_ of ∼110-115 Å, although this species could not be assigned to a defined intact FHC stoichiometry. Under the conditions tested, the FHC preparation also showed lower apparent ATPase and proteolytic activities than isolated FtsH. Together, these findings highlight the compositional and conformational heterogeneity of uncrosslinked FHC and reveal a more laterally splayed AAA+ arrangement in wild-type FtsH compared with the stabilized reference structure.

## Introduction

The membrane proteome of *E. coli* is maintained by a multilayered quality control network whose core proteolytic activity is performed by FtsH, an essential, ATP-dependent AAA+ metalloprotease anchored to the inner membrane via two N-terminal transmembrane helices ^1–3^. FtsH is unique among bacterial proteases in its capacity to extract and processively degrade both membrane-embedded and cytosolic substrates, coupling mechanical substrate dislocation which is powered by ATP hydrolysis, to zinc-dependent proteolysis within a single hexameric machine^1,4^.

FtsH degrades both soluble substrates, including σ32 (RpoH)^5^, CII^6^ and LpxC^7^, and integral membrane substrates such as unassembled SecY^8^. To keep this activity in check, the inner-membrane protein YccA modulates FtsH and protects complexes such as SecY from premature degradation^9^. Recently, Chai-Danino et al. (2026) showed that FtsH surveys the bilayer for membrane-exposed polar residues, a hallmark of misfolded or assembly-defective membrane proteins whose hydrophilic groups are displaced into the hydrophobic core; recognition of this degron depends on the FtsH transmembrane domain, placing the initial substrate decision within the membrane and before cytoplasmic unfolding^10^.

FtsH activity is further shaped by HflK and HflC, two single-pass inner-membrane proteins of the SPFH superfamily (stomatins, prohibitins, flotillins and HflKC)^11–13^. HflK and HflC form twelve heterodimers that oligomerize into a periplasmic scaffold encircling the transmembrane and periplasmic parts of FtsH. The resulting megadalton FtsH-HflKC holoenzyme (FHC) constitutes a membrane microdomain whose architecture has become a point of active debate^11,14,15^. Cross-linking mass spectrometry has additionally mapped extensive intra- and inter-protein contacts within recombinant FtsH-HflKC assemblies^16^.

Recent cryo-EM studies used chemical crosslinking (GraFix) during sample preparation and resolved a highly symmetric cage in which 24-subunit HflKC scaffold composed of 12 HflK-HflC heterodimers enclose four FtsH hexamers, with a twelve-fold periplasmic cap and a four-fold arrangement of hexamers inside^11,15^. On this basis HflKC was proposed to act as a static gate that sterically restricts substrate access to the FtsH pore^11,15^. It is worth noting that both cage studies already recorded particles with fewer than four hexamers, treated as partial or minority states (in one dataset, intact four-hexamer particles were under 10% of the population)^9^; so sub-stoichiometric occupancy was visible even under crosslinking, if not interpreted as such.

This picture was revised by Ghanbarpour et al. (2025)^17^. Their principal reconstruction (∼4.4 Å, locally 3.5 Å, no imposed symmetry) was obtained from DDM-solubilized complex, with glyco-diosgenin giving a similar lower-resolution map and detergent-free Carboxy-DIBMA/native-lipid preparations used at low resolution as a control. They described an asymmetric, nautilus-like HflKC assembly with a lateral aperture that could admit membrane-embedded substrates. Classification of that dataset resolved scaffolds carrying one or two FtsH hexamers and scaffolds with no FtsH density, and did not recover closed or four-hexamer states under the preparations. The nautilus-like assembly is associated with inward membrane curvature that correlates with lipid-scramblase activity, repositioning HflKC from a passive inhibitor to an active participant in membrane re-modelling.

Iqbal and Ghanbarpour et al. (2026) then tied conformational state to physiology by engineering a disulfide-crosslinked HflKC locked in the closed geometry^18^. Cells expressing the locked-closed variant recovered poorly from aminoglycoside stress, and a doubly opened state was seen in native complexes from tobramycin-treated cells. These observations support the physiological relevance of HflKC conformational state and its remodeling during aminoglycoside stress^18,19^.

What has not been tested is whether this compositional and conformational plasticity survives in a distinct biochemical preparation: a coexpressed, uncrosslinked complex solubilized in LMNG rather than DDM or polymer nanodiscs, and analysed without a native bilayer. We address this here, processing the isolated FtsH motor and the HflKC scaffold data separately from the same sample, and complement the cryo-EM with SEC-SAXS and bulk activity measurements.

## Results

### ATPase and protease activities of isolated FtsH and FHC

The purified FHC preparation contained HflK, HflC, and FtsH, as confirmed by SDS-PAGE (Fig. 1A; Supplementary Fig. S1A). For biochemical comparison, isolated FtsH and FHC were each assayed at a nominal concentration of 3 µM, with FHC molarity calculated on the basis of an assumed one-FtsH-hexamer/24-subunit-HflKC assembly (see Methods). Isolated FtsH showed higher apparent ATPase activity than the FHC preparation, with values of 9.97 U/mg and 3.85 U/mg, respectively, at 10 mM ATP (Fig. 1B). Across the ATP concentration series, phosphate release increased more strongly for isolated FtsH than for FHC.

**Figure 1.**
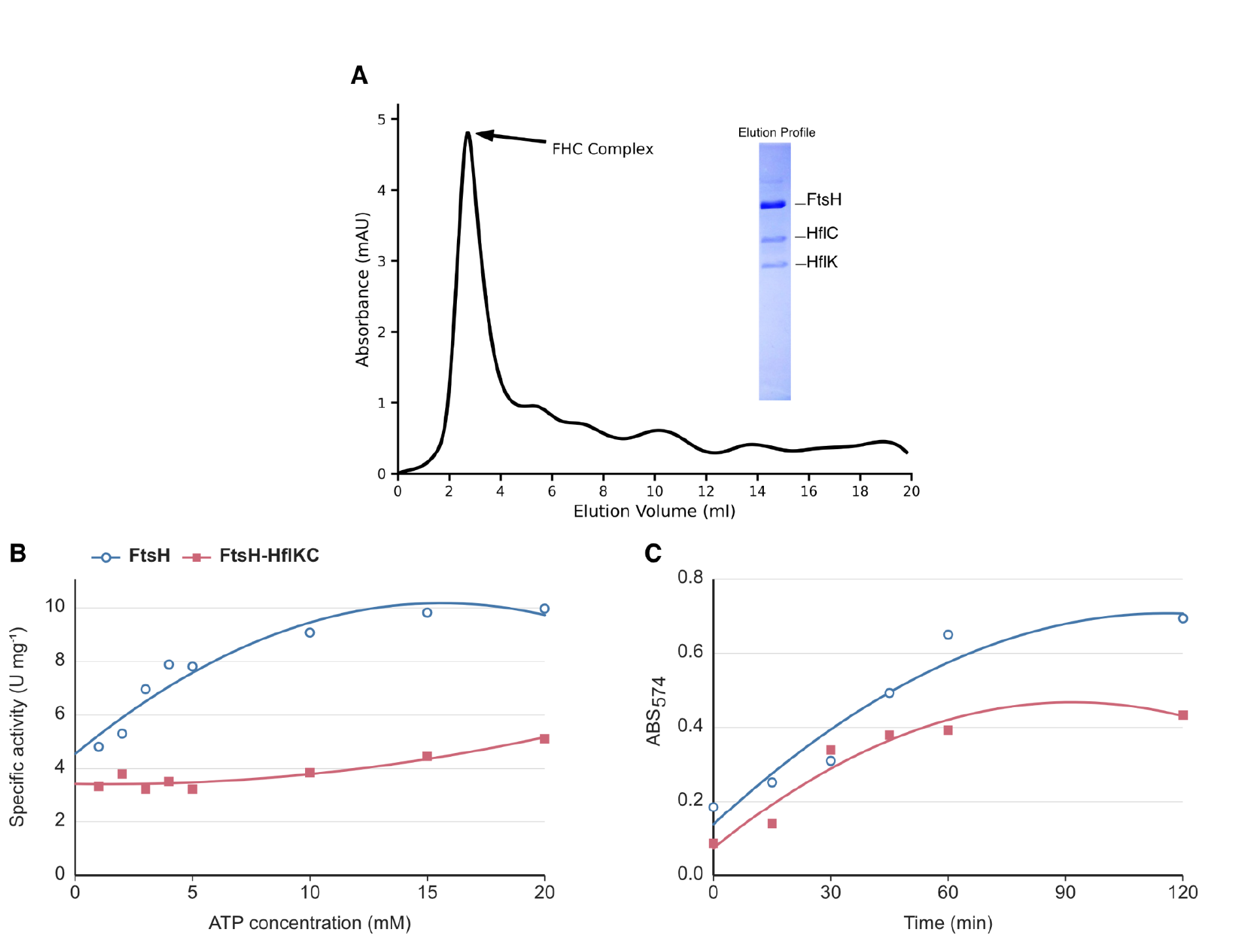
Purification and biochemical characterization of *E. coli* FtsH and FHC. (A) SEC elution profile of LMNG-solubilized FHC. The inset shows SDS-PAGE analysis of the eluted sample, confirming co-purification of HflK, HflC, and FtsH. (B) ATPase activity of isolated FtsH (blue) and FtsH-HflKC (rose), each assayed at 3 µM, over the indicated ATP concentration range. (C) Time course of resorufin-labeled casein cleavage by isolated FtsH (blue) and FtsH– HflKC (rose), each at a nominal concentration of 3 µM, monitored by absorbance at 574 nm. ATPase and protease measurements were obtained from a single experimental series (n = 1).

Proteolytic activity was assessed using resorufin-labeled casein as a general substrate. At 37°C, with the activity of FtsH in the absence of ATP defined as 100%, FHC showed relative activities of 39% without ATP and 40% in the presence of 5 mM ATP, whereas FtsH showed 91% relative activity in the presence of ATP (Fig. 1C). Thus, under the conditions tested, the FHC showed lower apparent ATPase and proteolytic activities than isolated FtsH. These measurements were obtained from a single experimental series and are interpreted descriptively.

### SEC-SAXS characterization of the LMNG-solubilized FHC preparation

To characterize the solution state of the LMNG-solubilized preparation independently of cryo-EM grid preparation, we performed SEC-SAXS. The CHROMIXS elution profile contained two overlapping peaks (Fig. 2A), and the later-eluting fraction was selected for analysis. The corresponding pair-distance distribution was broad and asymmetric, with an estimated *D_max_* of ∼110-115 Å (Supplementary Fig. S2B), and DAMMIF modelling yielded a compact, anisotropic envelope (Fig. 2B). The SAXS derived *D_max_* is smaller than the ∼140 Å height of the full-length wild-type FtsH reconstructed by cryo-EM in this study (Fig. 4B), indicating that the selected later-eluting species cannot be assigned directly to a defined intact FtsH-HflKC assembly. The earlier-eluting peak is compatible with a larger solution species; however, this fraction could not be processed further and therefore could not be structurally assigned.We therefore interpret the SAXS data at the level of the selected solution fraction. The contribution of LMNG to the measured scattering further limits direct interpretation of the envelope as a protein shape^20^.

**Figure 2.**
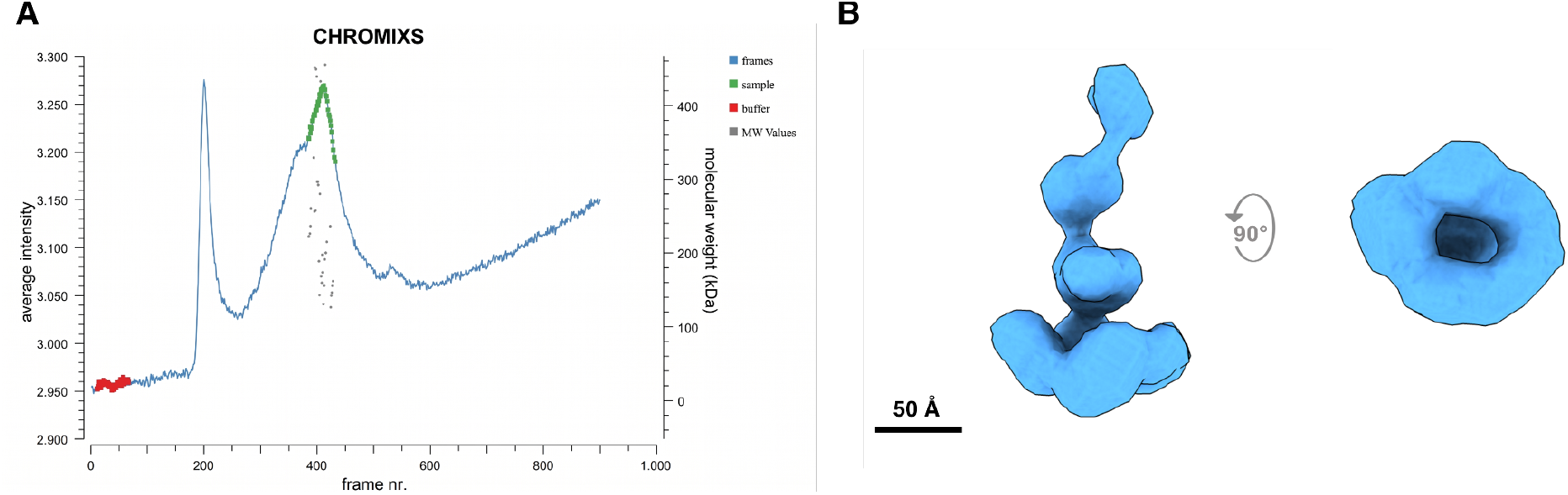
Solution characterization of *E. coli* FHC. (A) CHROMIXS profile of LMNG-solubilized FHC, showing two elution peaks. (B) *Ab initio* SAXS envelope (DAMMIF bead model) from the second SEC-SAXS peak fraction, shown in two orthogonal views.

### Cryo-EM captures HflKC scaffolds with variable FtsH occupancy

Negative-stain TEM showed dispersed particles in the purified preparation prior to cryo-EM analysis (Supplementary Fig. S1B). Single-particle data from LMNG-solubilized FHC gave 6,332 movies. After patch motion correction and CTF estimation, blob-based autopicking (100-600 Å diameter) returned 7,544,345 candidate particles, of which 2,853,018 were kept after inspection and initial 2D classification (Supplementary Fig. S3). The 2D class averages indicated substantial heterogeneity. 2D averages ranged from compact, pseudo-twelve-fold assemblies to asymmetric, nautilus-like arrangements, in line with prior native preparations^17^. We therefore split the data into two branches: isolated FtsH (Fig. 3A) and HflKC-containing assemblies (Fig. 3B,C), and processed them separately. Following iterative 2D classification and further particle selection, 11,866 FtsH particles and 21,442 HflKC-containing particles were retained for the final reconstruction branches.

**Figure 3.**
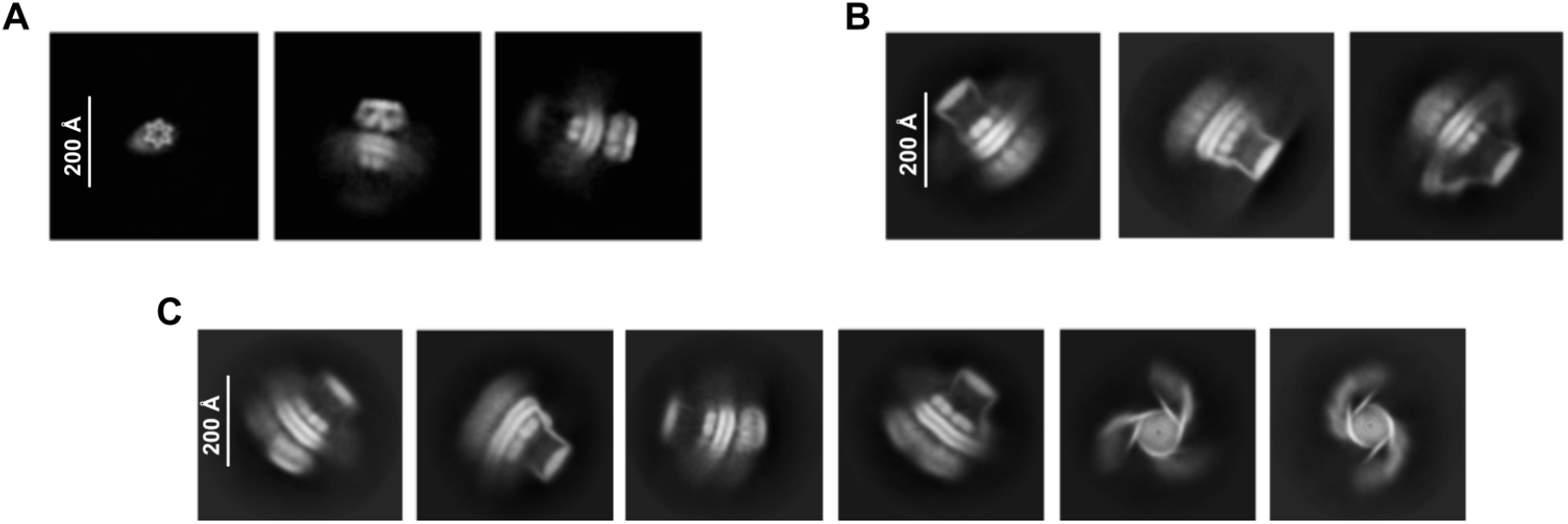
Stoichiometric variability of FtsH and FHC revealed by 2D classification. Representative 2D class averages of LMNG-solubilized FHC particles showing (A) isolated FtsH hexamers in top and side views, (B) HflKC scaffold associated with single FtsH hexamer, (C) HflKC scaffold accommodating two FtsH hexamers.

Within the HflKC branch, 2D class averages contained scaffolds with a single FtsH hexamer, and with two FtsH hexamers, within the same FtsH-HflKC preparation (Fig. 3B,C; Supplementary Fig. S4). Among the selected 2D classes assigned to these occupancy states, 9,793 particles were associated with one-FtsH classes and 8,329 with two-FtsH classes. These counts reflect the selected 2D classes and were not used to estimate the relative abundance of the two states. Multi-class 3D classification was subsequently attempted to separate the one- and two-FtsH occupancy states. However, the particle numbers available for the individual occupancy subclasses were insufficient to support stable, interpretable 3D reconstructions. The HflKC-containing particles were therefore used to obtain a consensus scaffold reconstruction, and the assignment of the one- and two-FtsH occupancy states remains based on the 2D class averages rather than on independently resolved 3D maps (Supplementary Fig. S3). This limitation applies specifically to FtsH occupancy within HflKC-containing assemblies; isolated FtsH particles formed a separate processing branch and yielded independent 3D reconstructions. The coexistence of HflKC-containing classes associated with one or two FtsH hexamers in the same uncrosslinked LMNG dataset extends the variable FtsH occupancy reported by Ghanbarpour et al. to a distinct biochemical preparation. These classes are not consistent with an obligatory four-hexamer cage as the sole assembly state in this preparation.

### A focused reconstruction of the isolated FtsH hexamer

From the FtsH branch, *ab initio* reconstruction and 3D classification yielded a full-length map, from which domain-focused masks were generated for separate refinement of the cytoplasmic and transmembrane/periplasmic regions (Supplementary Fig. S3). With C6 symmetry imposed, the cytoplasmic domain (CD) reached 5.32 Å resolution, resolving the overall AAA+ ring organization and secondary structure level features (Fig. 4E). The periplasmic/transmembrane domain (PD-TM) reconstruction reached 7.51 Å, while the full-length PD-TM-CD reconstruction reached 6.51 Å (Fig. 4B). All resolutions were determined by mask-corrected gold-standard Fourier Shell Correlation at the 0.143 criterion (Supplementary Fig. S5). Local resolution analysis showed the highest resolution within the cytoplasmic region (∼4-5 Å), with lower local resolution toward the transmembrane region (>10 Å) (Fig. 4C,D). The full-length FtsH hexamer spans approximately 140 Å, comprising an ∼80 Å cytoplasmic region, an ∼45 Å transmembrane region, and an ∼18 Å periplasmic domain, consistent with the overall dimensions reported for FtsH structures previously^21^.

**Figure 4.**
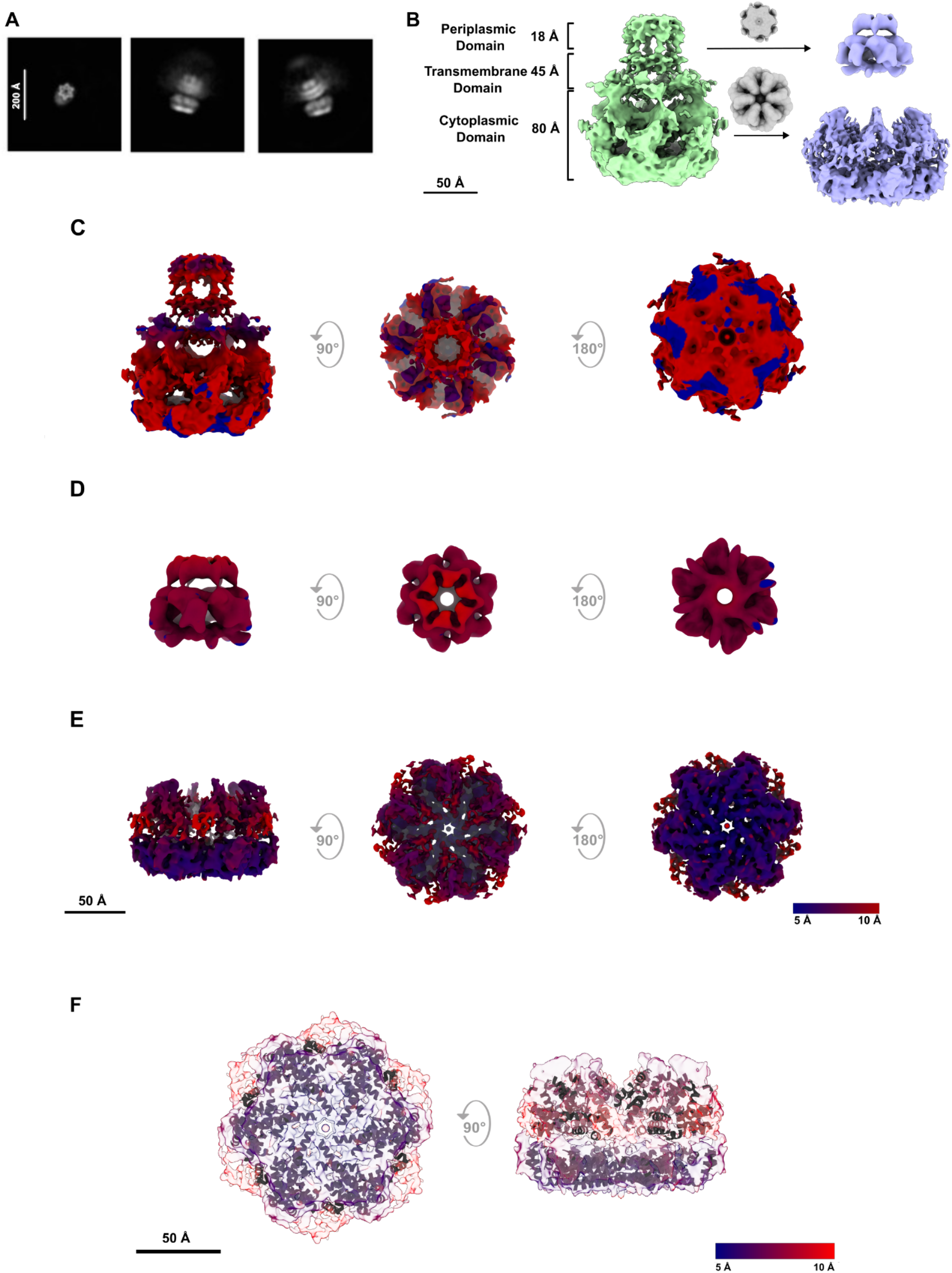
Cryo-EM structure of the isolated *E. coli* FtsH hexamer. (A) Representative 2D class averages of LMNG-solubilized FtsH. Top views (left) reveal hexameric features, while side views (right) resolve the cytosolic AAA+ domain. (B) 3D reconstruction of the FtsH imposed with C6 symmetry, left is a combination of separately processed FtsH-PD-TM and FtsH-CD structures in light purple whereas right is jointly processed FtsH-PD-TM-CD in light green. Local resolution estimation of the FtsH-PD-TM-CD (C) periplasmic and transmembrane domain map (D) and focused FtsH-PD-TM (side, top, bottom) map and (E) focused FtsH-CD map. The maps are colored by local resolution. (F) A fitted FtsH-CD AlphaFold model (AF-P0AAI3) is overlaid as a simplified secondary structure trace for orientation only.

The organization of the periplasmic domain was further examined by comparison with the CCW and rotated-helix conformation (RHC) maps identified in our previous study ^22^(Supplementary Fig. S6). The same periplasmic region was isolated from the sharpened and unsharpened TM-PD reconstructions, and the reference maps were fitted as rigid bodies in ChimeraX. To reduce the resolution difference between the reference maps, the higher-resolution CCW map was Gaussian-smoothed (σ = 0.95 Å) before fitting. A mildly smoothed version of the sharpened current density (σ = 0.5 Å) was also evaluated. For the sharpened map, the RHC reference gave a correlation coefficient of 0.4335, compared with 0.4070 for CCW. Mild smoothing retained the same relationship, with correlations of 0.4462 for RHC and 0.4217 for CCW. In the unsharpened map, the two references fitted similarly, with correlations of 0.8084 for RHC and 0.8195 for CCW. Thus, both previously described conformations are compatible with the present periplasmic domain density, and the modest change in relative fit across map representations does not support assignment to a unique CCW or RHC state.

### The wild-type, uncrosslinked FtsH-CD architecture

The 5.32 Å FtsH-CD reconstruction provides a structural view of wild-type, uncrosslinked FtsH without stabilizing substitutions or an added nucleotide analogue. The map retains the characteristic two tier organization of the cytosolic hexamer, with a lower protease ring and an upper AAA+ motor, but the two regions differ markedly in compactness^23–25^. The most prominent feature is the organization of AAA+ region, where the densities of six subunits are laterally displaced and more clearly separated, producing a broader, splayed arrangement with inter-subunit clefts. In contrast, the protease region forms a comparatively compact and continuous ring. Thus, the dominant structural variation within the wild-type FtsH-CD reconstruction is concentrated in the AAA+ motor region rather than representing a uniform expansion of the cytosolic assembly. To place this wild-type architecture in structural context, we compared it with EMD-32521/7WI4 (Fig. 5), a stabilized FtsH reconstruction obtained from the E252Q/E415Q double mutant in the presence of AMPPNP and following GraFix crosslinking (Supplementary Table S2). The reference retains a more tightly packed AAA+ domain in both the original and Gaussian-smoothed maps, whereas the overall envelope of the protease tier is similar between the reconstructions. The comparison therefore identifies the upper AAA+ motor as the principal region of architectural difference between the wild-type, uncrosslinked FtsH-CD map and the stabilized reference. At 5.32 Å resolution, this difference is resolved at the level of domain organization and overall compactness rather than individual side-chain rearrangements.

**Figure 5.**
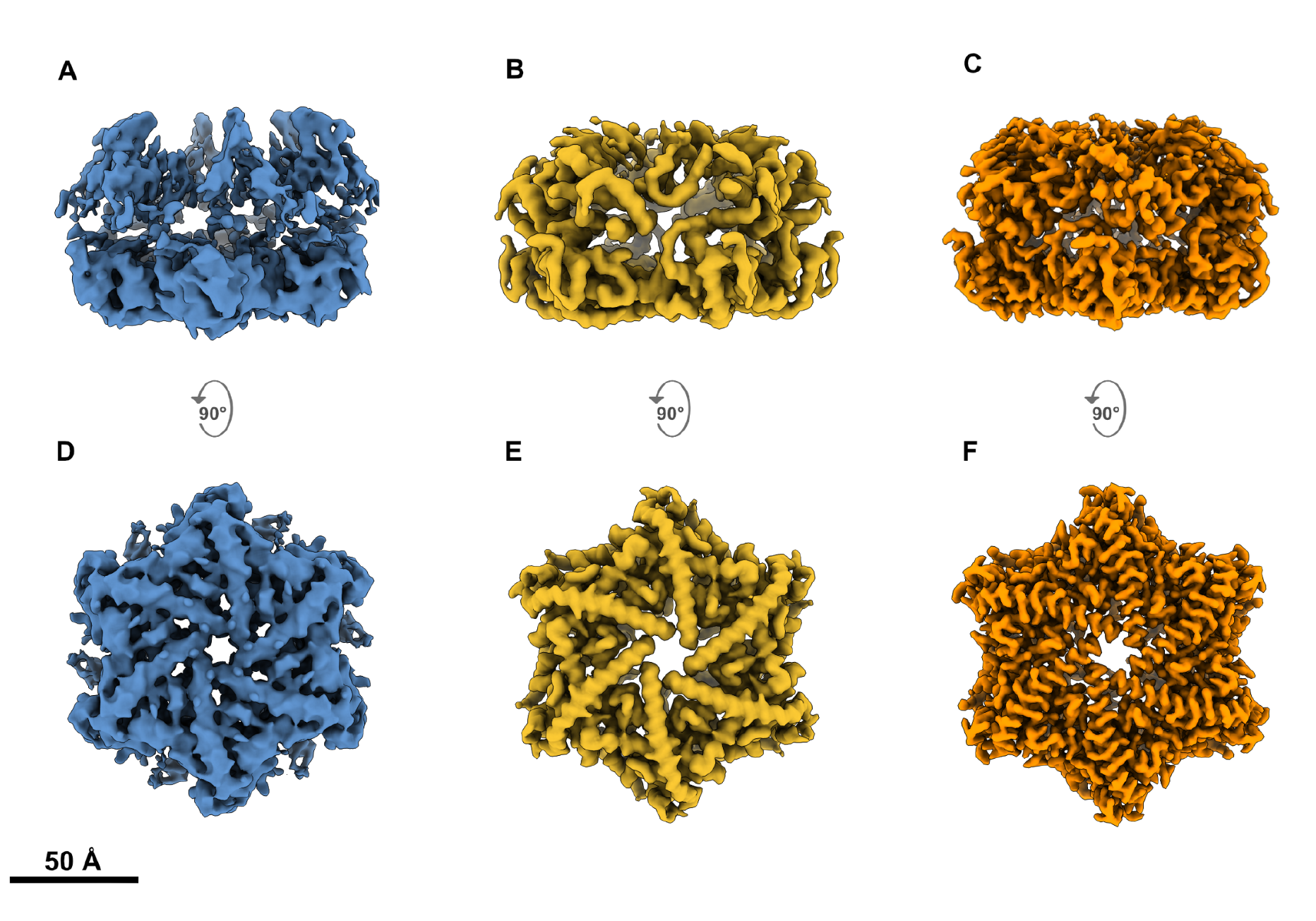
Map-level comparison of the wild-type FtsH-CD reconstruction with the EMD-32521/7WI4 map. (A) Side view of the sharpened wild-type FtsH-CD reconstruction at 5.32 Å. (B) Corresponding side view of the EMD-32521 reference map after Gaussian smoothing (σ = 2.0 Å) and alignment to the wild-type reconstruction. (C) Side view of the aligned EMD-32521 map at 3.4 Å. (D) Bottom view of the wild-type FtsH-CD reconstruction. (E) Corresponding bottom view of the Gaussian-smoothed EMD-32521 reference map. (F) Bottom view of the EMD-32521 map. Maps were displayed in a common orientation and contoured to yield comparable overall molecular envelopes. The wild-type reconstruction shows a less compact AAA+ region, whereas the lower protease domain remains similar between the reconstructions. The Gaussian-smoothed reference is included as a visual control for the effect of map sharpness.

### 3D Variability Analysis reveals conformational variability of HflKC scaffold

The HflKC particle stack (21,442 particles) exhibited conformational heterogeneity that limited the resolution of the consensus reconstruction to 9.63 Å. To characterize the dominant conformational variability within this ensemble, we applied 3D Variability Analysis (3DVA)^26^ in CryoSPARC. Although the HflKC particles were obtained from a C12 non-uniform refinement branch, 3DVA was performed in C1 without imposing additional symmetry. The dominant 3DVA component captured a coordinated change in the organization of the periplasmic scaffold (Fig. 6; Supplementary Movie S1). Representative intermediate volumes along this component span laterally splayed open-like geometries, a more compact and constricted geometry and more open-like geometries toward the opposite end of the variability coordinate (Fig. 6B,C). These intermediate volumes describe the range of conformational variability represented by the 3DVA component and should not be interpreted as time-resolved sequence of opening and closing events.

**Figure 6.**
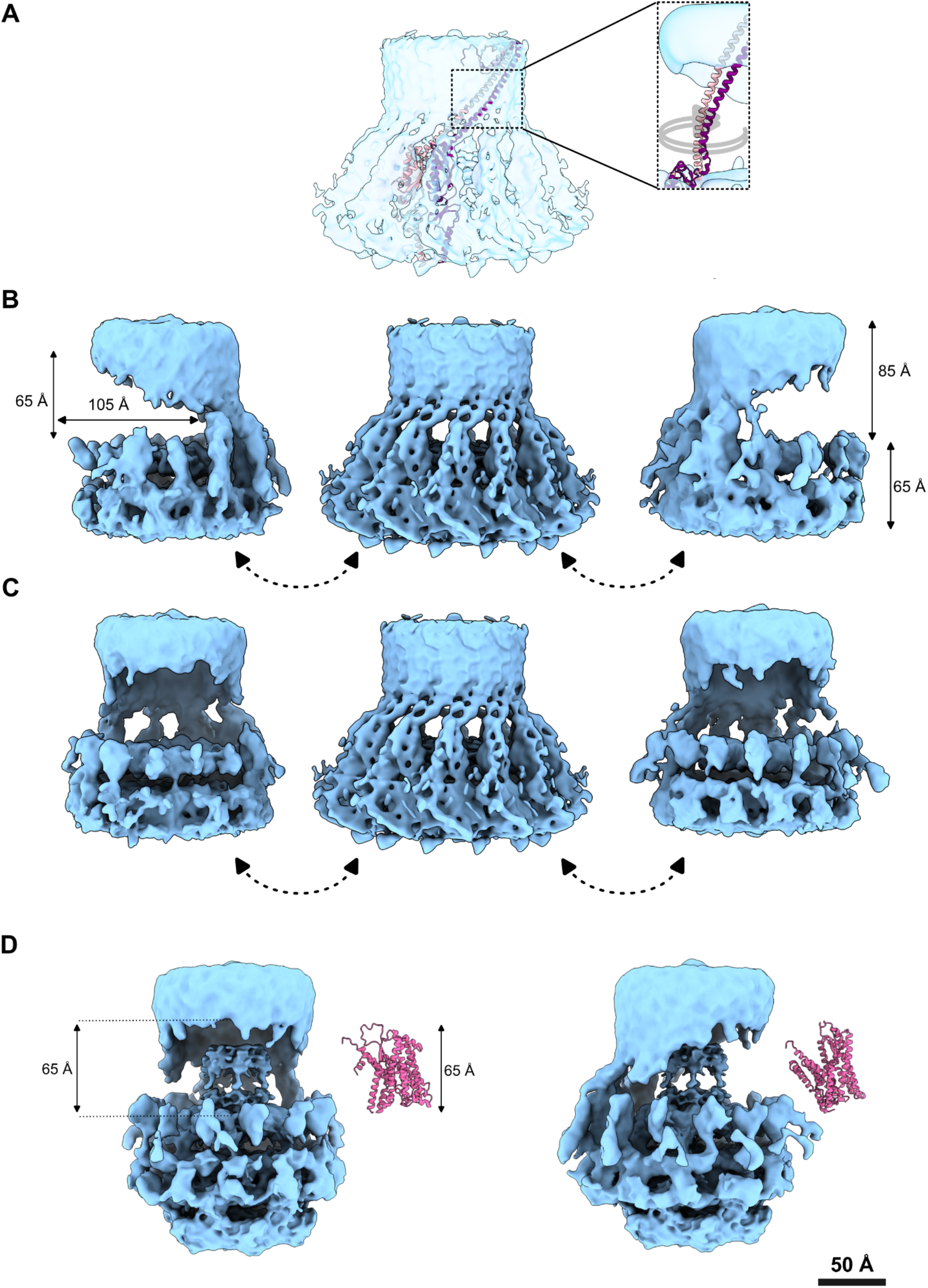
Conformational variability of the HflKC scaffold resolved by 3DVA. (A) Atomic models of isolated HflK and HflC monomers from PDB 7WI3 were rigid-body fitted into the HflKC map and further fitted to the open-gate conformation (inset). The open-gate map shown in the inset was low-pass filtered for clarity. (B, C) Three representative intermediate volumes sampled along the dominant 3DVA component illustrate the range of scaffold geometries in side and front views. The left and right volumes display open-like geometries with laterally splayed periplasmic domains, whereas the central volume is more constricted. Their ordering represents positions along the variability coordinate rather than a temporal sequence (see Supplementary Movie S1). (D) Front and side views of the 1:24 (FtsH hexamer:HflKC subunit) FHC showing the HflKC cavity with the known substrate SecY (PDB 6ITC) positioned next to the complex.

Notably, substantial scaffold variability is retained in the uncrosslinked, LMNG-solubilized preparation, demonstrating that chemical crosslinking is not required for HflKC to adopt distinct scaffold geometries. This observation supports an inherent capacity of the HflKC assembly to access multiple conformational states. Because the complex was analyzed in detergent and in the absence of a native lipid bilayer, however, the extent to which membrane lipids, curvature, or other native interactions modulate this conformational landscape cannot be determined from the present data. The open-like geometries captured by 3DVA are broadly consistent with the asymmetric architecture reported by Ghanbarpour et al. (2025) under detergent-free conditions, whereas the more constricted geometry resembles the occluded organization observed in stabilized cage structures. Together, these comparisons suggest that the conformational range sampled by the uncrosslinked HflKC preparation encompasses scaffold geometries related to both previously described architectural extremes. At the present resolution, however, residue level rearrangements at the FHC interface cannot be resolved. In the stabilized cage structure, residues K61/D62/S63 in the FtsH β2-β3 turn loop contact HflK β4; whether these contacts are retained or rearranged among the scaffold geometries observed here remains unresolved.

## Discussion

A central question concerning the FHC is whether HflKC constitutes a fixed inhibitory architecture or a structurally adaptable scaffold that accommodates different FtsH-associated states. Here, analysis of an uncrosslinked LMNG-solubilized preparation reveals heterogeneity at several levels, including variable FtsH occupancy, conformational variability of the HflKC scaffold, and structural differences between wild-type isolated FtsH and a previously stabilized FtsH reference. Biochemical measurements further show that the FHC preparation displays activity profiles distinct from isolated FtsH. Together, these observations support a view of FHC as a structurally heterogeneous assembly rather than a single fixed stoichiometric and conformational state.

### Stoichiometric variability challenges the fixed-valence cage model

The symmetric 4:24 (FtsH hexamer:HflKC subunit) cage resolved under crosslinking conditions^11,15^ implies a fixed binding geometry in which four FtsH hexamer positions are simultaneously and equivalently occupied. The coexistence of these classes indicates that more than one FtsH occupancy state can be accommodated within the same preparation. Our observation of HflKC-containing classes associated with one or two FtsH hexamers within the same crosslink-free dataset challenges an obligatory fixed-valence model. The coexistence of one- and two-FtsH-associated 2D classes demonstrates compositional heterogeneity within this preparation, but the selected-class counts do not establish relative state populations or *in vivo* stoichiometry.

### Solution-state heterogeneity observed by SEC-SAXS

SEC-SAXS revealed two overlapping elution features, indicating that the LMNG-solubilized preparation contains more than one solution-state species. The better-resolved later-eluting fraction yielded a compact anisotropic scattering envelope; however, its estimated *D_max_* of ∼110-115 Å is smaller than the ∼140 Å height of the full-length FtsH reconstruction obtained by cryo-EM. The selected SAXS species therefore cannot be assigned directly to an intact FHC complex or to the one- and two-FtsH classes observed by cryo-EM. Thus, the SAXS data provide independent evidence for heterogeneity within the biochemical preparation, but do not resolve the stoichiometry of the individual solution species.

### Conformational variability of the HflKC scaffold

The dominant C1 3DVA component spans open-like, more constricted, and again more open-like scaffold geometries, demonstrating substantial conformational variability within the HflKC particle ensemble. These geometries provide a structural context for the markedly different closed and asymmetric HflKC architectures reported previously, although the 3DVA series should not be interpreted as a temporal sequence connecting those structures. Because the 3DVA input particles were derived from a C12-refined HflKC branch, we interpret the resulting C1 3DVA series conservatively as a scaffold-level description of the dominant mode of scaffold variability, rather than as an atomically resolved asymmetric transition.

### Structural features of wild-type, uncrosslinked FtsH

In addition to the HflKC-containing assemblies, the cryo-EM dataset yielded a distinct population of isolated FtsH particles, providing a structural view of wild-type, uncrosslinked FtsH under the LMNG conditions used here. The FtsH-PD-TM reconstruction places the periplasmic domain within the conformational range identified in our previous analysis. Rigid-body comparisons with the previously described CCW and RHC states did not consistently favor one reference across map representations. The sharpened and mildly smoothed maps showed slightly higher agreement with RHC, whereas the unsharpened map showed a small preference for CCW. These differences were modest, indicating that the present reconstruction captures a periplasmic domain organization compatible with the structural landscape defined by both states rather than a uniquely resolved CCW- or RHC-like conformation. Importantly, the present map captures this periplasmic region together with the transmembrane domain in wild-type, uncrosslinked FtsH.

The most prominent feature of the 5.32 Å cytosolic-domain reconstruction is the reduced compactness of the AAA+ domain, where the subunit densities adopt a broader and more laterally splayed arrangement, while the protease domain remains comparatively compact. This suggests that the dominant structural variation is concentrated in the organization of the AAA+ motor rather than reflecting a uniform expansion of the entire cytosolic assembly. Comparison with EMD-32521/7WI4 places this architecture in context: the stabilized reference retains a more tightly packed AAA+ domain, including after Gaussian smoothing, whereas the lower protease domain remains similar between the two reconstructions. Because 7WI4 was obtained from an E252Q/E415Q double mutant in the AMPPNP-bound state following GraFix crosslinking, while the present map represents wild-type FtsH in an undefined nucleotide state without crosslinking, the observed difference cannot be attributed to any single biochemical or experimental factor. Nevertheless, the present wild-type, uncrosslinked FtsH reconstruction captures a less compact AAA+ organization than the stabilized reference. At the present resolution, this difference is most appropriately interpreted at the level of domain organization and conformational compactness rather than residue level rearrangements.

### HflKC association is accompanied by FtsH activity *in vitro*

Under the conditions tested, the FHC preparation showed lower apparent ATPase and proteolytic activities than isolated FtsH. HflKC has long been recognized as a modulator of FtsH proteolysis: early biochemical studies showed effects on the degradation of both membrane and soluble substrates, while subsequent work demonstrated that the direction and magnitude of HflKC-dependent regulation depend on the substrate and degradation pathway^6,9,13^. More recent structural and proteomic studies likewise argue against a simple constitutive inhibitory role. The closed HflKC architecture reported by Qiao et al. was proposed to restrict access of membrane substrates to FtsH^15^, whereas the asymmetric assemblies described by Ghanbarpour et al. contain an open entryway and were associated with both positive and negative effects on FtsH-dependent protein turnover^17^.

The lower caseinolytic and ATPase readouts observed here are consistent with this context-dependent modulation, but do not identify a specific regulatory mechanism. Casein is a soluble general protease substrate and therefore does not directly report membrane-substrate access, while ATPase and proteolytic outputs are not necessarily coupled proportionally^27^. We therefore interpret the present measurements as showing that the FHC preparation has a biochemical profile distinct from isolated FtsH under the conditions tested.

### FHC as a structurally heterogeneous assembly

Taken together, the structural observations support a model in which FHC can populate more than one organizational state rather than adopting a single obligatory architecture. Variable FtsH occupancy within HflKC-containing classes, conformational variability of the HflKC scaffold, and the distinct architecture observed for isolated wild-type FtsH all point to substantial structural plasticity in the uncrosslinked LMNG-solubilized preparation. This view is broadly consistent with the asymmetric and variable-occupancy assemblies reported in recent studies and contrasts with a model in which the symmetric four-hexamer cage represents the sole FHC state. The lower apparent ATPase and proteolytic activities observed for the FHC preparation indicate a distinct biochemical profile relative to isolated FtsH under the conditions tested.

### Limitations and future directions

The present study does not resolve the one- and two-FtsH HflKC-containing states as independent 3D reconstructions, and the 9.63 Å HflKC consensus limits interpretation to the scaffold level. The biochemical measurements were obtained from a single experimental series, while interpretation of the SEC-SAXS fractions is limited by detergent scattering and uncertain particle composition. Higher-resolution analysis of defined FHC states, replicated biochemical measurements, and reconstitution in a membrane-like environment will be required to establish how the observed structural heterogeneity relates to FtsH regulation.

## Conclusion

We show that an uncrosslinked, LMNG-solubilized FtsH-HflKC contains HflKC-associated 2D classes with one or two FtsH hexamers and displays substantial scaffold-level conformational variability. Independently, isolated FtsH particles yielded a 5.32 Å wild-type FtsH-CD reconstruction with a less compact AAA+ arrangement than the stabilized reference, although the experimental differences between these reconstructions preclude attribution to a single factor. Together, these observations extend recent evidence for FtsH-HflKC structural heterogeneity to a distinct biochemical preparation and argue against the symmetric four-hexamer cage as the sole organizational state represented under these conditions.

## Material and Methods

### Construction of FtsH-HflKC Plasmid

Construction of the FHC plasmid (pAce-HflKC-His10_x_pDS-FtsH-3strep) was performed using the ACEMBL MultiColi modular expression system^28^. 10 ng of plasmids were transformed into the *E. coli* 10-beta cells (NEB, UK) by heat shock (42°C, 30 sec), incubated in LB at 37°C for 1 h, and plated on LB agar with 100 μg/ml ampicillin and spectinomycin.

### Overexpression, Solubilization and Purification of FtsH-HflKC and FtsH

FHC plasmid was transformed into *E. coli* Lemo21(DE3) competent cells for protein overexpression. A 20 mL aliquot of this overnight culture was used to inoculate 1 L of 2xYT medium for large-scale expression. Induction was initiated with 400 μM IPTG and 0.2% arabinose when the culture reached an OD_600_ of 0.3-0.6. Following incubation for 4 hours at 30 °C, cells were harvested by centrifugation (Beckman Coulter Avanti JLA-8.1000 rotor, 5,000 rpm, 25 min, 4°C). The resulting pellet was resuspended in breaking buffer (50 mM Tris-HCl, pH 7.5, 100 mM NaCl) and disrupted by sonication (VibraCell VCX130, Sonics, USA) using 3 cycles of 1-minute pulses (1 s on, 1 s off) at 50% amplitude. Lysates were clarified by centrifugation. Membrane pellets (∼0.7 g per condition) were solubilized using 1% (w/w) Lauryl Maltose Neopentyl Glycol (LMNG) detergent and incubated at 4°C for 1 hour in breaking buffer (50 mM Tris-HCl, pH 7.5, 100 mM NaCl) with gentle agitation using a rotator. After solubilization, samples were centrifuged at 100,000 × g for 45 min (Beckman Optima XE, Ti45 rotor), and supernatants were analyzed by SDS-PAGE (12%). The detergent-solubilized supernatant containing FHC was subjected to Strep-tag affinity chromatography according to the manufacturer’s protocol (IBA LifeSciences, US). The solubilized protein was loaded onto the affinity column and incubated for 10 min, after which the flow-through was collected. The column was washed with Buffer W (50 mM Tris-HCl, pH 7.5, 100 mM NaCl) containing 0.005% LMNG and elution was performed with 2.5 mM desthiobiotin in Buffer E (50 mM Tris-HCl, pH 7.5, 100 mM NaCl, 2.5 mM desthiobiotin).

Isolated Strep-tagged *E.coli* FtsH used for biochemical comparison was purified using the same membrane isolation, detergent solubilization, and Strep-affinity procedure as described above for the FtsH-HflKC preparation. Following LMNG solubilization, insoluble material was removed by centrifugation at 100,000 × *g* for 45 min. The solubilized fraction was loaded onto Strep affinity resin and incubated for 10 min before collection of the flow-through. The resin was washed with 50 mM Tris-HCl (pH 7.5), 100 mM NaCl containing 0.005% LMNG, and bound FtsH was eluted in the same buffer supplemented with 2.5 mM desthiobiotin.

### Size-exclusion chromatography

Following Strep-affinity purification, FHC eluate was concentrated using 100-kDa molecular-weight-cutoff centrifugal concentrators (Merck, DE) and subsequently purified by size-exclusion chromatography using a HiLoad 16/600 Superdex 200 column. The column was equilibrated in 50 mM Tris-HCl (pH 7.5), 100 mM NaCl and 0.0025% LMNG, corresponding to the affinity-purification buffer without desthiobiotin and with the detergent concentration reduced by half. Size-exclusion chromatography was performed at a flow rate of 1.0 mL/min. Fractions corresponding to the major elution peak were pooled and concentrated using 100-kDa MWCO centrifugal concentrators.

### Biochemical activity assays

ATPase activity of purified FHC and FtsH was measured using the EnzChek Phosphate Assay Kit (Thermo Fisher Scientific, US), which detects inorganic phosphate P_i_ released upon ATP hydrolysis through the purine nucleoside phosphorylase (PNP) dependent conversion of MESG. A phosphate calibration curve was generated using KH_2_PO_4_ standards containing 0, 10, 20, 40, 60, 80, and 100 µM inorganic phosphate, and absorbance was measured at 360 nm. ATPase reactions were prepared in a total volume of 600 µL containing 30 µL of 20× reaction buffer supplied with the kit, 120 µL MESG, 6 µL PNP (1 U), and either isolated FtsH or FHC at a final nominal concentration of 3 µM. For the FHC preparation, molar concentration was calculated assuming an assembled complex comprising one FtsH hexamer associated with the 24-subunit HflKC scaffold (12 HflK and 12 HflC subunits; approximate molecular mass, 1.4 MDa). Reaction mixtures were pre-incubated for 10 min at room temperature before addition of ATP. ATP concentrations of 1, 2, 3, 4, 5, 10, 15, and 20 mM were tested in the presence of 1 mM KCl and 1 mM NaCl, and phosphate release was monitored spectrophotometrically at 360 nm.

Proteolytic activity was measured using resorufin-labeled casein (Sigma-Aldrich, DE) as a general protease substrate. The substrate solution contained 0.4% resorufin-labeled casein in ddH_2_O, the incubation buffer contained 0.2 M Tris-HCl (pH 7.8) and 0.02 M CaCl_2_, the stop solution contained 5% (w/v) trichloroacetic acid, and the activity buffer contained 0.5 M Tris-HCl (pH 8.8). Each reaction contained 30 µL substrate solution, 30 µL incubation buffer, and 60 µL of FtsH or FHC adjusted to a nominal concentration of 3 µM, using the same concentration basis described above. Blank reactions contained 60 µL ddH_2_O in place of protein. Proteolysis reactions were incubated at 37°C for 15, 30, 45, 60, and 120 min. Where indicated, 5 mM ATP was included. Reactions were stopped by addition of 288 µL of 5 % trichloroacetic acid and incubated for a further 10 min at 37 °C. Samples were centrifuged at 12,000 × g for 5 min, after which 240 µL of the supernatant was mixed with 360 µL activity buffer and absorbance was measured immediately at 574 nm. Relative proteolytic activities were calculated by normalizing each condition to the activity of isolated FtsH measured at 37 °C in the absence of ATP, which was defined as 100%.

### Negative-staining

FtsH-HflKC samples were prepared separately at 0.05 mg/ml concentrations containing LMNG. To investigate homogeneity, carbon-coated grids (CF200 TEM Grids, 200 Mesh, Cu) were incubated in 70% ethanol for one hour and the grids were allowed to dry. A 5 µl sample was placed on the grid and left for one minute. The excess sample was then blotted off the grid with blotting paper and the grid was placed on 20 µl of ddH_2_O, left for 15 seconds and blotted again with filter paper. Then the grid was finally placed in 15 µl 2% Uranyl Acetate, left for one minute and the excess was removed with blotting paper. The samples were examined using FEI Tecnai G2 S-Twin at Ankara University, Faculty of Arts and Sciences, Department of Biology.

### Sample preparation and cryo-EM data collection

3 µL aliquot of FHC at 0.5 mg/mL was applied to a Quantifoil R1.2/1.3 carbon grid (Agar Scientific) that had been glow-discharged at 30 mA for 1 min. Grids were plunge-frozen using a Vitrobot Mark IV (Thermo Fisher Scientific) at 16 °C and 80% humidity with a blotting time of 1.6 s. A total of 6,332 movies, each comprising 52 frames, were collected using EPU on a 300 kV Titan Krios microscope equipped with a Falcon 4 detector at the Thermo Fisher Scientific NanoPort facility (Eindhoven, Netherlands). Data were acquired at a nominal magnification of 130,000× with a total exposure of 50 e⁻/Å² and a defocus range of −0.8 to −2.0 µm. The pixel size was 0.929 Å.

### Cryo-EM image processing

A total of 6,332 movies, each containing 52 frames, were processed in CryoSPARC^29^. The movies were subjected to patch-based motion correction followed by contrast-transfer-function estimation using Patch CTF. Initial particle picking was performed using the blob picker with particle diameters ranging from 100 to 600 Å, yielding 7,544,345 candidate particles. Following particle inspection and removal of false-positive picks, 2,853,018 particles were retained for further classification. Iterative 2D classification revealed substantial particle heterogeneity and enabled separation of the dataset into two principal processing branches corresponding to isolated FtsH and HflKC-containing assemblies.

The isolated FtsH branch was further subjected to iterative 2D classification, followed by *ab initio* reconstruction in C1 and 3D classification. The 11,866 particle class was selected for subsequent refinement and subjected to an initial non-uniform refinement with C6 symmetry. From this particle set, the full-length FtsH-PD-TM-CD reconstruction was generated using Flex Reconstruction, yielding a final resolution of 6.51 Å. Focused refinements using masks encompassing the cytosolic and transmembrane/periplasmic regions yielded the FtsH-CD and FtsH-PD-TM reconstructions at 5.32 Å and 7.51 Å, respectively. For visualization purposes, a copy of the focused FtsH-CD reconstruction was post-processed using DeepEMhancer^30^. This post-processing was used only for figure visualization and was not used for particle refinement or FSC-based resolution determination. The reported 5.32 Å resolution was determined from the mask-corrected gold-standard FSC at the 0.143 criterion.

The HflKC-containing branch was further curated by iterative 2D classification, yielding a final stack of 21,442 particles. These particles were subjected to *ab initio* reconstruction in C1 followed by non-uniform refinement with C12 symmetry. A subsequent masked C12 non-uniform refinement yielded the 9.63 Å HflKC consensus reconstruction. The HflKC particle stack was further analyzed by 3D Variability Analysis in C1 without imposing additional symmetry. All reported resolutions were determined using the mask-corrected gold-standard FSC at the 0.143 criterion^31^ (Supplementary Fig. S5 and Supplementary Table S1).

### Map-level comparison of the FtsH cytosolic and periplasmic domains

The reference cryo-EM map EMD-32521, corresponding to the FtsH reconstruction associated with PDB 7WI4, was obtained from the EMDB and compared with the 5.32 Å wild-type FtsH-CD reconstruction generated in this study. The maps were aligned by rigid-body map-to-map fitting in UCSF ChimeraX using the `fitmap` command, with the wild-type FtsH-CD reconstruction used as the reference density^32^. To evaluate whether differences in map sharpness contributed to the apparent difference in AAA+ tier compactness, the higher-resolution EMD-32521 reference was additionally Gaussian-smoothed using σ = 2.0 Å. The Gaussian width was selected empirically as a deliberately stringent smoothing control to attenuate high-spatial-frequency detail in the reference while retaining the overall cytosolic domain envelope; it was not intended to match the nominal resolution of the two reconstructions. The original and Gaussian-smoothed reference maps were compared with the wild-type FtsH-CD reconstruction in identical side and bottom orientations. Structural interpretation was restricted to differences in overall domain organization and compactness that remained apparent after smoothing.

The periplasmic domain region was cropped using an identical spatial region from the sharpened and unsharpened FtsH-PD-TM reconstructions. The previously reported CCW (EMD-66269) and RHC (EMD-66013) reference maps were compared with the current density by rigid-body map-to-map fitting in UCSF ChimeraX. Because the CCW reference contained substantially finer density features than the current reconstruction, it was Gaussian-smoothed using σ = 0.95 Å as a moderate smoothing control before fitting. The sharpened current map was additionally evaluated after mild Gaussian smoothing (σ = 0.5 Å) to test the sensitivity of the comparison to map sharpening. These smoothing parameters were selected empirically to reduce differences in high-frequency map detail while preserving the overall periplasmic-domain envelope, rather than to impose a common nominal resolution. Map correlations were calculated after rigid-body fitting using the ChimeraX `fitmap` correlation.

### 3D Variability Analysis

To characterize continuous conformational heterogeneity of the HflKC scaffold, the HflKC particle stack containing 21,442 particles was subjected to 3D Variability Analysis in CryoSPARC. The particles were obtained from the HflKC processing branch after C12 non-uniform refinement; however, 3DVA itself was performed in C1 without imposing additional symmetry. Intermediate volume series were generated using the “intermediates” output mode, with volumes downsampled to a box size of 256 pixels and filtered to 11.5 Å for visualization. Three variability components were examined, and the dominant component describing the largest scaffold-level conformational variation was used for Fig. 6 and Supplementary Movie S1. The ordering of the intermediate volumes reflects their positions along the 3DVA coordinate and should not be interpreted as a temporal opening–closing sequence.

### Sample Preparation for SAXS and Data Processing

The purified FHC (1.5 mg/ml) was transported to the European Molecular Biology Laboratory (EMBL, Hamburg), BioSAXS, PETRA-III P12 for Small Angle X-Ray Scattering analysis^33^. Here, 100 µl of sample was further purified using the HPLC Superdex 200 Increase 5/150 GL column in order to obtain homogeneity. SAXS data were collected over a *q* range of 0.007–0.4 Å⁻¹. SEC-SAXS data were processed using the ATSAS software package^34^. The elution profile was inspected using CHROMIXS^35^, and the later-eluting fraction was selected for further analysis. The pair-distance distribution function, *p(r)*, and the corresponding *D_max_* were obtained within ATSAS, and *ab initio* bead modelling was performed using DAMMIF.

## Acknowledgments

This study was supported by the Scientific and Technological Council of Türkiye (TÜBİTAK) (Project No: 118C225). G.G., and B.V.K. were funded by TÜBİTAK BİDEB 2232 International Outstanding Researchers Program (Project No: 118C225). G.G. was also funded by TÜBİTAK 2211-A National Graduate Fellowship Program. C.S. acknowledges funding by a BBSRC Responsive Mode Grant (BB/P000940/1) and a Wellcome Trust Investigator Grant (210701/Z/18/Z). SAXS data were collected at the EMBL Hamburg BioSAXS beamline (P12) at PETRA III (HH-SAXS-1278), operated by DESY, and we acknowledge the support and hospitality of the DESY staff. We thank the Turkish Energy, Nuclear and Mining Research Authority (TENMAK) for funding under the “Support to SESAME Activities” scheme. Furthermore, we would like to thank Dr. Hakan Eskizengin for his assistance with ns-TEM analysis. We also acknowledge Thermo Fisher Scientific (Eindhoven, Netherlands) for providing access to cryo-EM facilities and infrastructure.

## Data Availability

The cryo-EM reconstructions generated in this study are being prepared for deposition in the Electron Microscopy Data Bank (EMDB). Accession numbers are not yet available and will be included in an updated version of this preprint once assigned. Additional data supporting the findings of this study are presented in the manuscript and its Supplementary Information.

## Declaration of AI-assisted tools

During the preparation of this manuscript, the author used ChatGPT (OpenAI, accessed September 2026) to improve the structure of the draft and to suggest alternative titles for subsections.

## References

(1) Ito, K.; Akiyama, Y. Cellular Functions, Mechanism of Action, and Regulation of FtsH Protease. Annual Review of Microbiology 2005, 59, 211–231. 10.1146/annurev.micro.59.030804.121316

(2) Bittner, L.-M.; Arends, J.; Narberhaus, F. When, How and Why? Regulated Proteolysis by the Essential FtsH Protease in Escherichia Coli. Biological Chemistry 2017, 398 (5–6), 625–635. doi:10.1515/hsz-2016-0302

(3) Langklotz, S.; Baumann, U.; Narberhaus, F. Structure and Function of the Bacterial AAA Protease FtsH. Biochim Biophys Acta 2012, 1823 (1), 40–48. 10.1016/j.bbamcr.2011.08.015

(4) Chiba, S.; Akiyama, Y.; Ito, K. Membrane Protein Degradation by FtsH Can Be Initiated from Either End. J Bacteriol 2002, 184 (17), 4775–4782. 10.1128/JB.184.17.4775-4782.2002

(5) Tomoyasu, T.; Gamer, J.; Bukau, B.; Kanemori, M.; Mori, H.; Rutman, A. J.; Oppenheim, A. B.; Yura, T.; Yamanaka, K.; Niki, H. Escherichia Coli FtsH Is a Membrane-Bound, ATP-Dependent Protease Which Degrades the Heat-Shock Transcription Factor Sigma 32. EMBO J 1995, 14 (11), 2551–2560. 10.1002/j.1460-2075.1995.tb07253.x

(6) Kihara, A.; Akiyama, Y.; Ito, K. Host Regulation of Lysogenic Decision in Bacteriophage Lambda: Transmembrane Modulation of FtsH (HflB), the cII Degrading Protease, by HflKC (HflA). Proc Natl Acad Sci U S A 1997, 94 (11), 5544–5549. 10.1073/pnas.94.11.5544

(7) Führer, F.; Langklotz, S.; Narberhaus, F. The C-Terminal End of LpxC Is Required for Degradation by the FtsH Protease. Mol Microbiol 2006, 59 (3), 1025–1036. 10.1111/j.1365-2958.2005.04994.x

(8) Akiyama, Y.; Kihara, A.; Tokuda, H.; Ito, K. FtsH (HflB) Is an ATP-Dependent Protease Selectively Acting on SecY and Some Other Membrane Proteins *. Journal of Biological Chemistry 1996, 271 (49), 31196–31201. 10.1074/jbc.271.49.31196

(9) Kihara, A.; Akiyama, Y.; Ito, K. Different Pathways for Protein Degradation by the FtsH/HflKC Membrane-Embedded Protease Complex: An Implication from the Interference by a Mutant Form of a New Substrate Protein, YccA; 1998.

(10) Chai-Danino, M.; Ravensary-Modin, N.; Vladimirov, V. I.; Onyshchuk, T.; Plöhn, M.; Barshap, A. B. D.; Bsoul, A.; Peled-Zehavi, H.; Fluman, N. Membrane-Embedded Polar Residues Target Membrane Proteins for Degradation by the Quality Control Protease FtsH. Nature Communications 2026, 17 (1), 3067. 10.1038/s41467-026-69829-8

(11) Ma, C.; Wang, C.; Luo, D.; Yan, L.; Yang, W.; Li, N.; Gao, N. Structural Insights into the Membrane Microdomain Organization by SPFH Family Proteins. Cell Research 2022, 32 (2), 176–189. 10.1038/s41422-021-00598-3

(12) Yokoyama, H.; Matsui, I. The Lipid Raft Markers Stomatin, Prohibitin, Flotillin, and HflK/C (SPFH)-Domain Proteins Form an Operon with NfeD Proteins and Function with Apolar Polyisoprenoid Lipids. Crit Rev Microbiol 2020, 46 (1), 38–48. 10.1080/1040841X.2020.1716682

(13) Kihara, A.; Akiyama, Y.; Ito, K. A Protease Complex in the Escherichia Coli Plasma Membrane: HflKC (HflA) Forms a Complex with FtsH (HflB), Regulating Its Proteolytic Activity against SecY. EMBO J 1996, 15 (22), 6122–6131.

(14) Saikawa, N.; Akiyama, Y.; Ito, K. FtsH Exists as an Exceptionally Large Complex Containing HflKC in the Plasma Membrane of Escherichia Coli. In Journal of Structural Biology; 2004; Vol. 146, pp 123–129. 10.1016/j.jsb.2003.09.020

(15) Qiao, Z.; Yokoyama, T.; Yan, X.-F.; Beh, I. T.; Shi, J.; Basak, S.; Akiyama, Y.; Gao, Y.-G. Cryo-EM Structure of the Entire FtsH-HflKC AAA Protease Complex. Cell Reports 2022, 39 (9), 110890. 10.1016/j.celrep.2022.110890

(16) Akkulak, H.; İnce, H. K.; Goc, G.; Lebrilla, C. B.; Kabasakal, B. V.; Ozcan, S. Structural Proteomics of a Bacterial Mega Membrane Protein Complex: FtsH-HflK-HflC. Int J Biol Macromol 2024, 269 (Pt 1), 131923. 10.1016/j.ijbiomac.2024.131923

(17) Ghanbarpour, A.; Telusma, B.; Powell, B. M.; Zhang, J. J.; Bolstad, I.; Vargas, C.; Keller, S.; Baker, T. A.; Sauer, R. T.; Davis, J. H. An Asymmetric Nautilus-like HflK/C Assembly Controls FtsH Proteolysis of Membrane Proteins. EMBO J 2025, 44 (9), 2501–2513. 10.1038/s44318-025-00408-1

(18) Iqbal, N.; Keller, S.; Ghanbarpour, A. Structural Plasticity of the Membrane-Bound Protein Degradation Assembly Supports Bacterial Adaptation to Stress. Cell Reports 2026, 45 (4), 117231. 10.1016/j.celrep.2026.117231

(19) Wessel Aimee K.; Yoshii Yutaka; Reder Alexander; Boudjemaa Rym; Szczesna Magdalena; Betton Jean-Michel; Bernal-Bayard Joaquin; Beloin Christophe; Lopez Daniel; Völker Uwe; Ghigo Jean-Marc. Escherichia Coli SPFH Membrane Microdomain Proteins HflKC Contribute to Aminoglycoside and Oxidative Stress Tolerance. Microbiology Spectrum 2023, 11 (4), e01767–23. 10.1128/spectrum.01767-23

(20) Molodenskiy, D. S.; Mertens, H. D. T.; Svergun, D. I. An Automated Data Processing and Analysis Pipeline for Transmembrane Proteins in Detergent Solutions. Scientific Reports 2020, 10 (1), 8081. 10.1038/s41598-020-64933-1

(21) Carvalho, V.; Prabudiansyah, I.; Kovacik, L.; Chami, M.; Kieffer, R.; van der Valk, R.; de Lange, N.; Engel, A.; Aubin-Tam, M.-E. The Cytoplasmic Domain of the AAA+ Protease FtsH Is Tilted with Respect to the Membrane to Facilitate Substrate Entry. Journal of Biological Chemistry 2021, 296. 10.1074/jbc.RA120.014739

(22) Göc, G.; Yadav, S. K. N.; Orriss, G.; Borucu, U.; Berger, I.; Schaffitzel, C.; Kabasakal, B. V. Cryo-EM Structure of the FtsH Periplasmic Domain Reveals Functional Dynamics. ACS Chemical Biology 2026, 21 (4), 844–851. 10.1021/acschembio.5c01025

(23) Bieniossek, C.; Schalch, T.; Bumann, M.; Meister, M.; Meier, R.; Baumann, U. The Molecular Architecture of the Metalloprotease FtsH. Proc Natl Acad Sci U S A 2006, 103 (9), 3066–3071. 10.1073/pnas.0600031103

(24) Bieniossek, C.; Niederhauser, B.; Baumann, U. M. The Crystal Structure of Apo-FtsH Reveals Domain Movements Necessary for Substrate Unfolding and Translocation. Proc Natl Acad Sci U S A 2009, 106 (51), 21579–21584. 10.1073/pnas.0910708106

(25) Suno, R.; Niwa, H.; Tsuchiya, D.; Zhang, X.; Yoshida, M.; Morikawa, K. Structure of the Whole Cytosolic Region of ATP-Dependent Protease FtsH. Mol Cell 2006, 22 (5), 575–585. 10.1016/j.molcel.2006.04.020

(26) Punjani, A.; Fleet, D. J. 3D Variability Analysis: Resolving Continuous Flexibility and Discrete Heterogeneity from Single Particle Cryo-EM. J Struct Biol 2021, 213 (2), 107702. 10.1016/j.jsb.2021.107702

(27) Yang, Y.; Gunasekara, M.; Muhammednazaar, S.; Li, Z.; Hong, H. Proteolysis Mediated by the Membrane-Integrated ATP-Dependent Protease FtsH Has a Unique Nonlinear Dependence on ATP Hydrolysis Rates. Protein Sci 2019, 28 (7), 1262–1275. 10.1002/pro.3629

(28) Bieniossek, C.; Nie, Y.; Frey, D.; Olieric, N.; Schaffitzel, C.; Collinson, I.; Romier, C.; Berger, P.; Richmond, T. J.; Steinmetz, M. O.; Berger, I. Automated Unrestricted Multigene Recombineering for Multiprotein Complex Production. Nature Methods 2009, 6 (6), 447–450. 10.1038/nmeth.1326

(29) Punjani, A.; Rubinstein, J. L.; Fleet, D. J.; Brubaker, M. A. cryoSPARC: Algorithms for Rapid Unsupervised Cryo-EM Structure Determination. Nature Methods 2017, 14 (3), 290–296. 10.1038/nmeth.4169

(30) Sanchez-Garcia, R.; Gomez-Blanco, J.; Cuervo, A.; Carazo, J. M.; Sorzano, C. O. S.; Vargas, J. DeepEMhancer: A Deep Learning Solution for Cryo-EM Volume Post-Processing. Communications Biology 2021, 4 (1), 874. 10.1038/s42003-021-02399-1

(31) Rosenthal, P. B.; Henderson, R. Optimal Determination of Particle Orientation, Absolute Hand, and Contrast Loss in Single-Particle Electron Cryomicroscopy. J Mol Biol 2003, 333 (4), 721–745. 10.1016/j.jmb.2003.07.013

(32) Meng, E. C.; Goddard, T. D.; Pettersen, E. F.; Couch, G. S.; Pearson, Z. J.; Morris, J. H.; Ferrin, T. E. UCSF ChimeraX: Tools for Structure Building and Analysis. Protein Sci 2023, 32 (11), e4792. 10.1002/pro.4792

(33) Blanchet, C. E.; Spilotros, A.; Schwemmer, F.; Graewert, M. A.; Kikhney, A.; Jeffries, C. M.; Franke, D.; Mark, D.; Zengerle, R.; Cipriani, F.; Fiedler, S.; Roessle, M.; Svergun, D. I. Versatile Sample Environments and Automation for Biological Solution X-Ray Scattering Experiments at the P12 Beamline (PETRA III, DESY). J Appl Crystallogr 2015, 48 (Pt 2), 431–443. 10.1107/S160057671500254X

(34) Manalastas-Cantos, K.; Konarev, P. V.; Hajizadeh, N. R.; Kikhney, A. G.; Petoukhov, M. V.; Molodenskiy, D. S.; Panjkovich, A.; Mertens, H. D. T.; Gruzinov, A.; Borges, C.; Jeffries, C. M.; Svergun, D. I.; Franke, D. ATSAS 3.0: Expanded Functionality and New Tools for Small-Angle Scattering Data Analysis. J Appl Crystallogr 2021, 54 (Pt 1), 343–355. 10.1107/S1600576720013412

(35) Graewert, M. A.; Franke, D.; Jeffries, C. M.; Blanchet, C. E.; Ruskule, D.; Kuhle, K.; Flieger, A.; Schäfer, B.; Tartsch, B.; Meijers, R.; Svergun, D. I. Automated Pipeline for Purification, Biophysical and X-Ray Analysis of Biomacromolecular Solutions. Scientific Reports 2015, 5 (1), 10734. 10.1038/srep10734

