## Supporting Information for "Structural heterogeneity of uncrosslinked FtsH-HflKC assemblies includes variable FtsH occupancy and a distinct wild-type FtsH architecture"

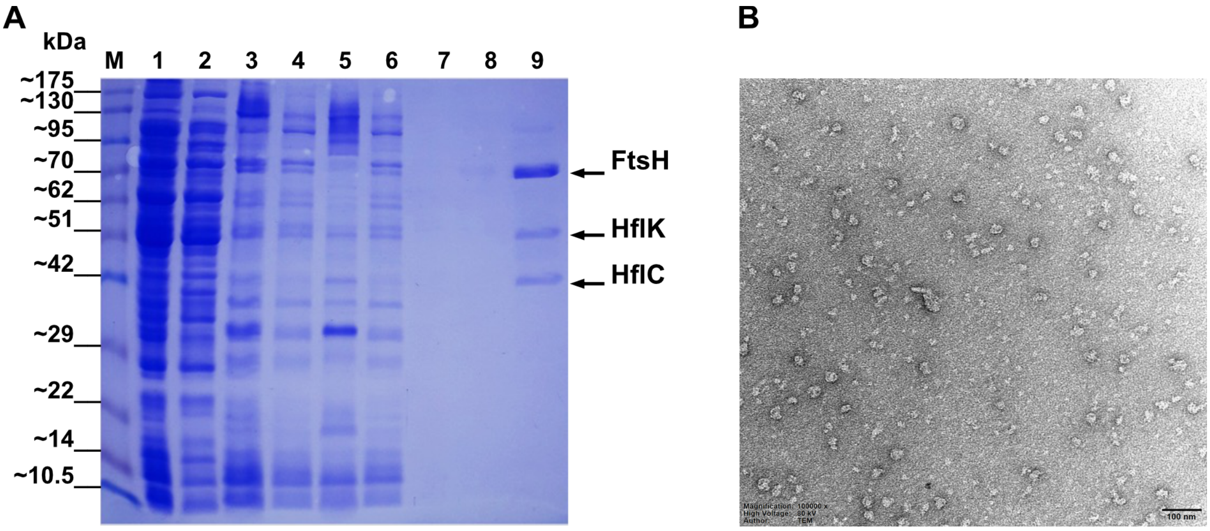


**Supplementary Figure S1.** Additional characterization of the purified FHC preparation. (A) Full SDS-PAGE image of the LMNG-solubilized FHC used for biochemical and structural characterization. Lane 1, cell lysate; lane 2, membrane fraction after LMNG incubation and before centrifugation; lane 3, LMNG-solubilization mixture before centrifugation; lane 4, membrane pellet remaining after solubilization; lane 5, flow-through; lane 6, wash fraction; lanes 7-9, elution fractions 1-3, respectively. The strongest FtsH-HflK-HflC bands were observed in elution fraction 3 (lane 9). The bands corresponding to FtsH, HflK, and HflC are indicated. (B) Representative negative-stain transmission electron micrograph of the purified

FHC, showing dispersed particles with heterogeneous particle dimensions and orientations. Scale bar, 100 nm.

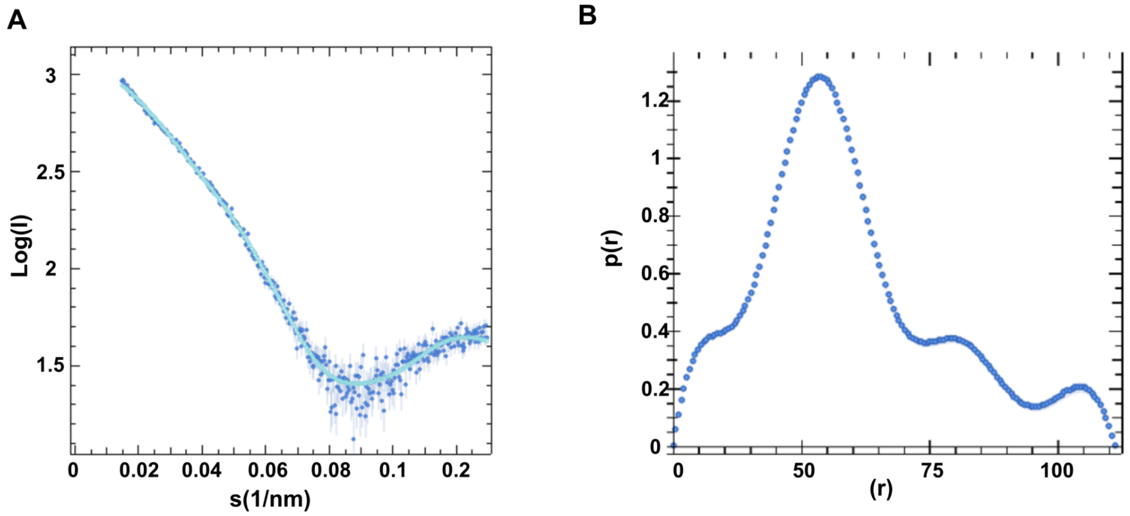


**Supplementary Figure S2.** SAXS characterization of the selected later-eluting SEC fraction of the LMNG-solubilized FHC. (A) SAXS scattering profile of the selected fraction plotted as log(I) against the scattering vector *q*. (B) Corresponding pair-distance distribution function, *p(r)*, showing a broad asymmetric distribution with an estimated $D_{max}$ of ~110-115 Å.

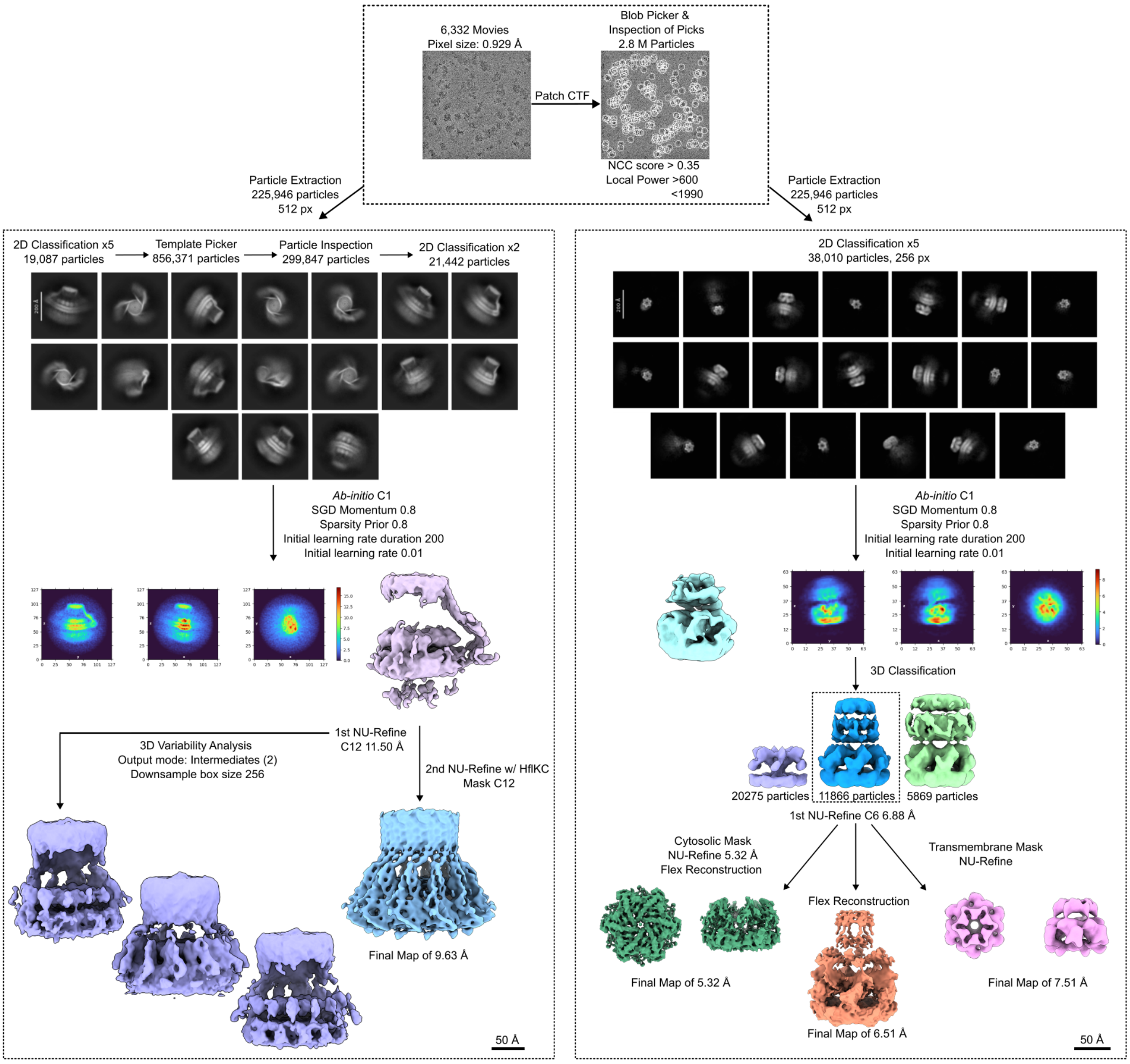


**Supplementary Figure S3.** CryoSPARC processing workflow used for the uncrosslinked FtsH-HflKC complex and isolated FtsH particles. After patch motion correction and CTF estimation of 6,332 movies, blob-based particle picking yielded 7,544,345 putative particles. Following particle inspection and iterative 2D classification, 2,853,018 particles were retained for further sorting. The dataset was then separated into two processing branches according to the dominant particle features observed in 2D class averages.

In the FHC/HflKC branch, 2D classification revealed HflKC-containing assemblies with variable FtsH occupancy. Multi-class 3D classification was attempted to separate the one- and two-FtsH occupancy states observed in the 2D class averages. However, the particle numbers available for the individual occupancy subclasses were insufficient to support reliable independent 3D reconstructions. Further subclass-specific refinement was therefore not pursued. Instead, the curated HflKC-containing particle stack was subjected to *ab initio* reconstruction and non-uniform refinement, yielding a consensus HflKC/FHC map from

21,442 particles at 9.63 Å resolution. To characterize continuous scaffold motion, particles from the C12 non-uniform refinement branch were further analyzed by 3D Variability Analysis in C1 using intermediate mode. Representative intermediate volumes from the dominant variability component illustrate open-like, more constricted, and more open-like scaffold geometries along the 3DVA coordinate. Their ordering reflects position along the variability component and should not be interpreted as a temporal sequence.

In the isolated FtsH branch, FtsH-containing particles were processed separately to reconstruct the full-length motor and its individual regions. Following *ab initio* reconstruction and 3D classification, the selected FtsH class was refined to generate the full-length FtsH map. Domain-focused masks derived from the full-length reconstruction were then used to refine the cytoplasmic domain and the transmembrane/periplasmic region separately. This workflow yielded reconstructions of full-length FtsH, focused FtsH-CD, and focused FtsH-TM-PD at 6.51 Å, 5.32 Å, and 7.51 Å resolution, respectively.

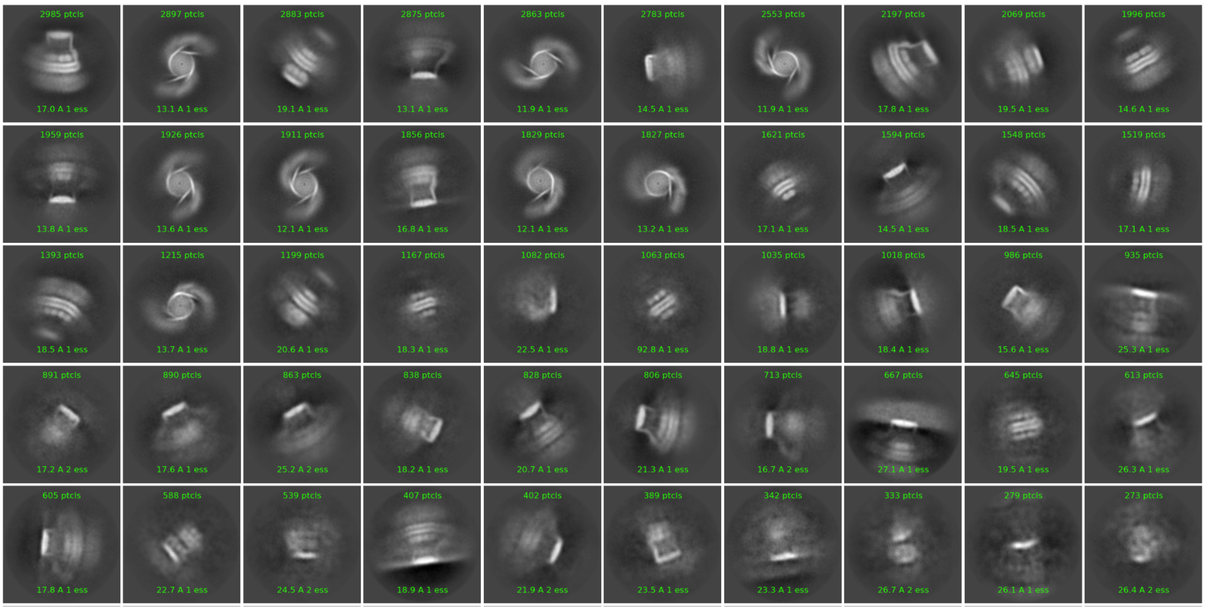

**Supplementary Figure S4.** CryoSPARC 2D classes of FHC particles. The numbers displayed on each class average indicate the number of particles assigned to that class by CryoSPARC. Classes corresponding to HflKC scaffolds associated with one or two FtsH hexamers were observed, supporting the coexistence of multiple FtsH:HflKC stoichiometries within the same dataset.

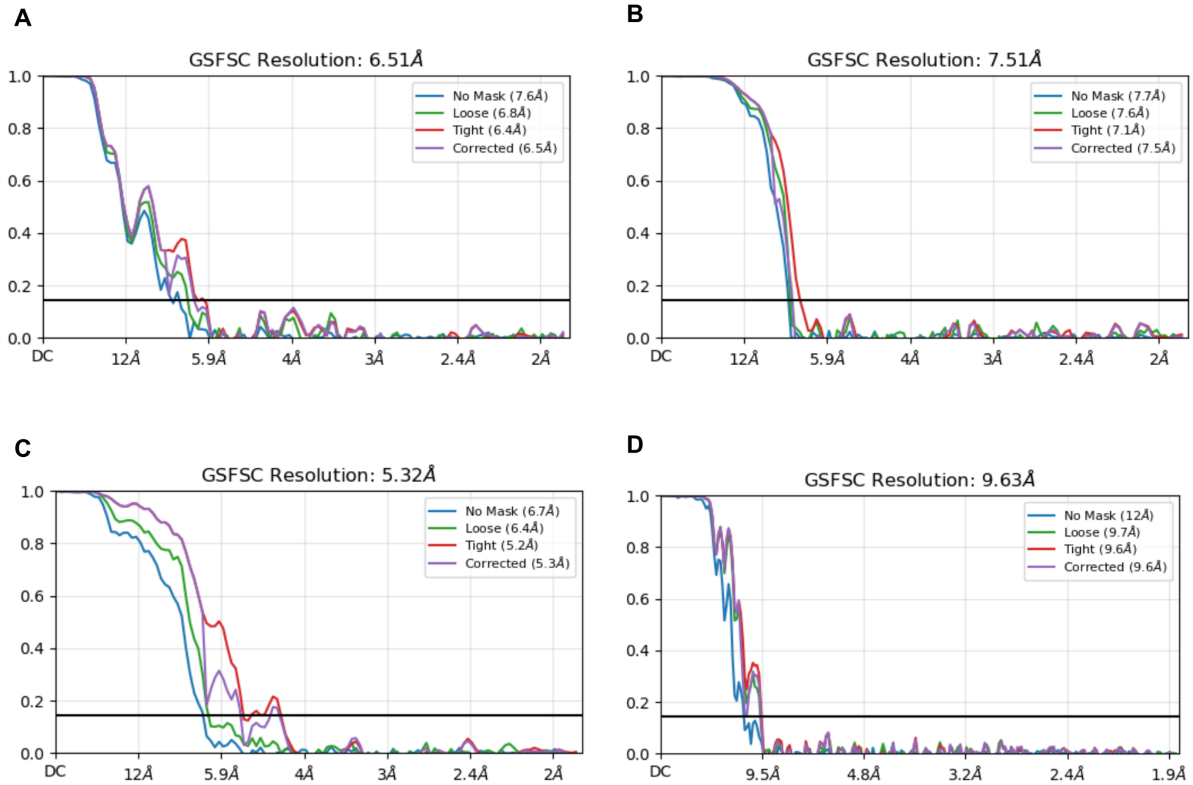


**Supplementary Figure S5.** Gold-standard FSC validation of the FtsH and HflKC reconstructions. GS-FSC curves for (A) flexible reconstruction map of FtsH-TM-PD-CD, (B) the focused FtsH-TM-PD, (C) the focused FtsH-CD, and (D) HflKC. The resolutions reported in the main text are based on the mask-corrected FSC at the 0.143 criterion and correspond to 6.51 Å, 7.51 Å, 5.32 Å, and 9.63 Å, respectively.

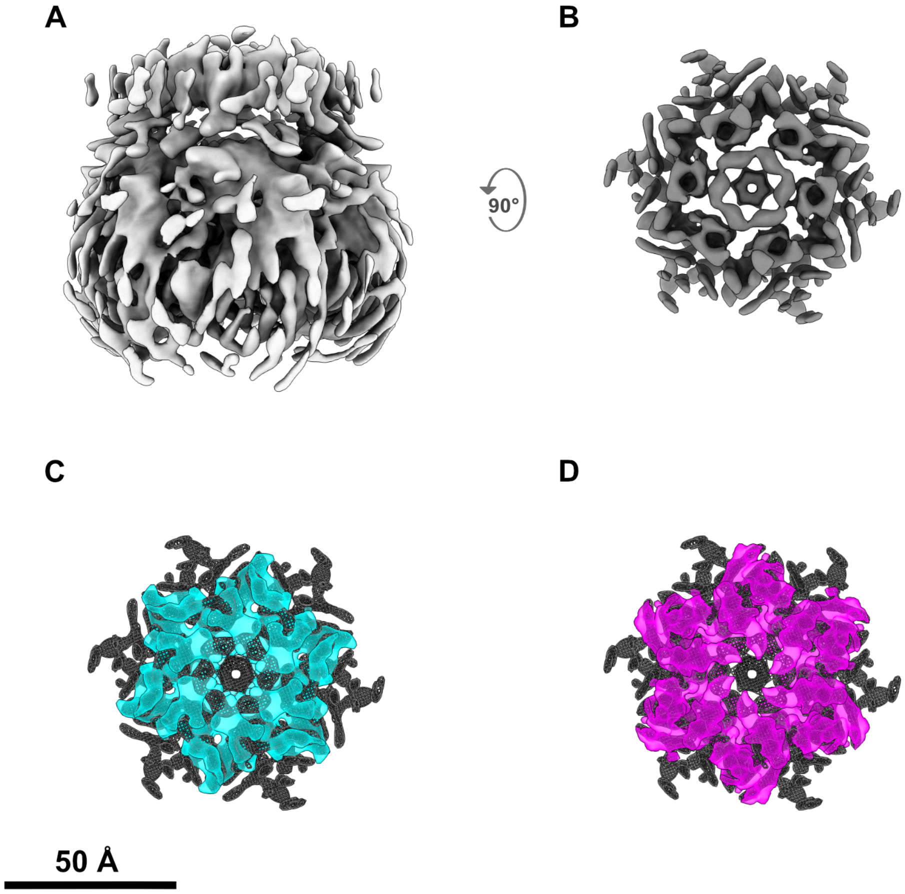


**Supplementary Figure S6.** Comparison of the FtsH periplasmic domain density with previously characterized CCW and RHC conformations. (A) Side view of the sharpened FtsH-PD-TM reconstruction used in the present study. (B) Top view of the cropped periplasmic domain density used for map level comparison. (C) Overlay of the sharpened periplasmic domain density (gray mesh) with the previously reported rotated-helix conformation (RHC; EMD-66013, cyan). (D) Equivalent overlay with the CCW reference (EMD-66269, magenta). Because the CCW reference was determined at higher resolution, it was Gaussian-smoothed (σ = 0.95 Å) before comparison. The RHC and CCW reference maps were fitted as rigid bodies to the same cropped periplasmic domain region in ChimeraX.

**Supplementary Table S1.**

| Feature | FtsH-TM-PD-CD | FtsH-TM-PD | FtsH-CD | FHC |
|---|---|---|---|---|
| Microscope | Titan Krios | | | |
| Detector | Falcon 4 | | | |
| Pixel size | 0.929 Å (binned by 2) | | | |

| | | | | |
|---|---|---|---|---|
| Nominal Magnification | 130000 | | | |
| Exposure ($e^-$/Å$^2$) | 50 $e^-$/Å$^2$ | | | |
| Defocus range | -0.8 to -2 µm | | | |
| Box size | 256 px | | | 512 px |
| Corrected gold-standard FSC (0.143) | 6.51 Å | 7.51 Å | 5.32 Å | 9.63 Å |
| Unmasked gold-standard FSC (0.143) | 7.6 Å | 7.7 Å | 6.7 Å | |
| Map sharpening | - | - | *DeepEMhancer* | - |
| Symmetry | C6 | | | C12* |

**C12 refers to the HflKC consensus non-uniform refinement; 3DVA was performed in C1.*

**Supplementary Table S2.** Constructs and experimental differences.

| Feature | This study, FtsH-CD | Qiao et al., 7WI4 |
|---|---|---|
| FtsH mutation | Wild type | E252Q/E415Q |
| Nucleotide state | No nucleotide analog added (wild-type enzyme); density at the ATPase site not assignable at ~5.3 Å | AMPPNP·$Mg^{2+}$ |
| Crosslinking | None | GraFix/glutaraldehyde |
| Processing | CryoSPARC focused mask | Relion subregion subtraction |
| Symmetry | C6 | C6 |
